# ProtNote: a multimodal method for protein-function annotation

**DOI:** 10.1101/2024.10.17.618952

**Authors:** Samir Char, Nathaniel Corley, Sarah Alamdari, Kevin K. Yang, Ava P. Amini

## Abstract

Understanding the protein sequence-function relationship is essential for advancing protein biology and engineering. However, fewer than 1% of known protein sequences have human-verified functions. While deep learning methods have demonstrated promise for protein function prediction, current models are limited to predicting only those functions on which they were trained. Here, we introduce ProtNote, a multimodal deep learning model that leverages free-form text to enable both supervised and zero-shot protein function prediction. ProtNote not only maintains near state-of-the-art performance for annotations in its train set, but also generalizes to unseen and novel functions in zero-shot test settings. We envision that ProtNote will enhance protein function discovery by enabling scientists to use free text inputs, without restriction to predefined labels – a necessary capability for navigating the dynamic landscape of protein biology.

## Introduction

Proteins, the building blocks of cellular life, exhibit astonishing functional diversity. Indeed, in domains as varied as medicine [1, 2], agriculture [3], and the food industry [4], scientists continue to discover novel and valuable applications for proteins. Before applying proteins to downstream tasks, practitioners must first understand their functions. However, high-fidelity functional annotations are sparse, with fewer than 1% of the sequence entries in the UniProt [5] database containing human-verified functions (Fig. 1A). Developing tools to predict protein function from sequence automatically is paramount to not only inform our biochemical understanding of proteins but also to accelerate applications to an expanding set of domains.

**Fig. 1.**
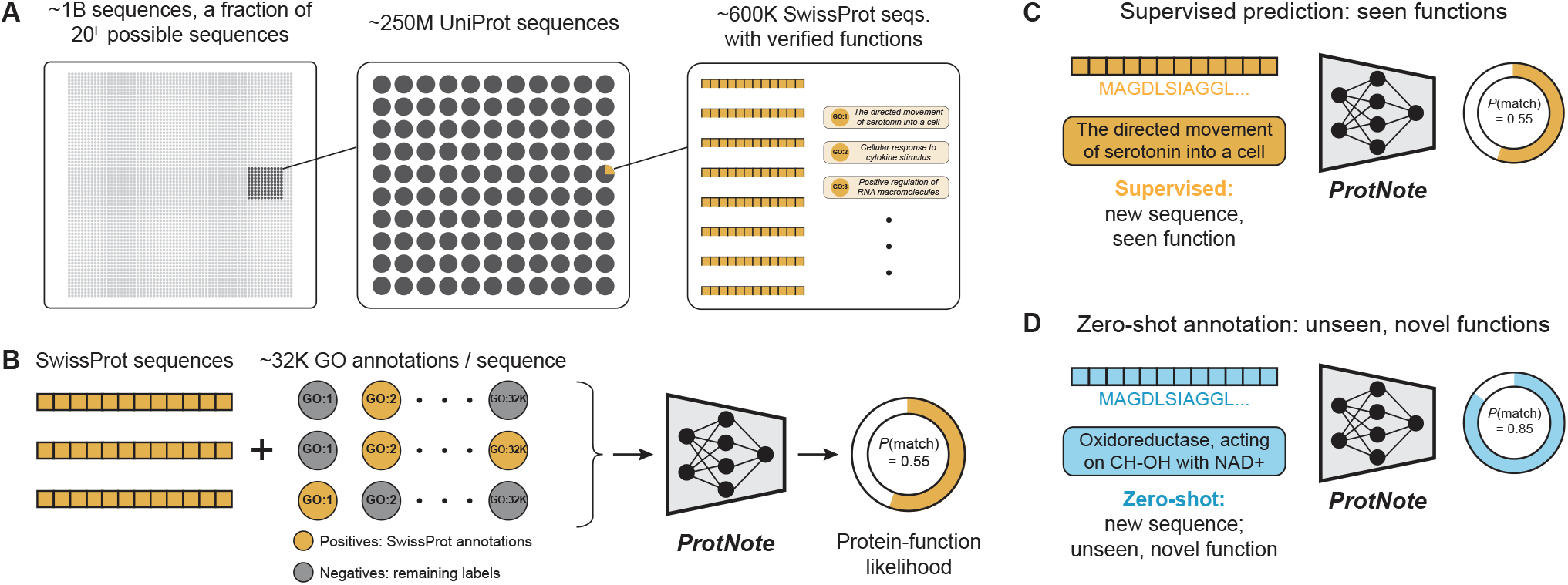
Protein function annotation with ProtNote. **(A)** A small fraction of all possible protein sequences are recorded in the UniProt database [5]. An even smaller portion, available in SwissProt [26], have human-verified functions. Text descriptions of function are assigned Gene Ontology term labels. **(B)** For a given sequence from SwissProt, positive labels (yellow) are applicable true SwissProt annotations; negative terms (grey) are the labels that do not apply to the sequence and are pulled from the Gene Ontology database [11, 12]. ProtNote is trained on amino acid sequences and the textual descriptions from the positive and negative GO terms and outputs a predicted protein-function likelihood for each input pair. **(C-D)** At inference, ProtNote can be deployed in both supervised (C) and zero-shot (D) settings. ProtNote takes unseen protein-function pairs as inputs and outputs the probability that the proteins perform the functions they were paired with, in both supervised prediction over seen functions (C, yellow) and zero-shot annotation over unseen, novel functions (D, blue).

Existing computational tools to predict functional annotations fall into two archetypes: homology-based methods and *de novo* methods. Homology-based methods encompass the majority of traditional strategies and include sequence alignment-based tools like Basic Local Alignment Search Tool (BLAST) [6] and family classification techniques like profile Hidden Markov Models [7, 8]. While effective and explainable, these methods are difficult to scale [9] and tend to underperform in remote homology situations, due to their reliance on database search via sequence similarity. On the other hand, *de novo* methods, most commonly machine learning models, do not directly rely on the query sequence’s homology and instead construct a representation of the protein from which to infer function. Notable recent *de novo* approaches include ProteInfer [9] and DeepGOPlus [10], both of which predict Gene Ontology (GO) [11, 12] annotations and Enzyme Commission (EC) numbers [13]; methods that specialize in EC number prediction, such as CLEAN and HiFi-NN [14, 15]; ProtENN [16], which infers protein families from the Pfam database [17]; and ProteinBERT [18], a GO-annotation-aware protein language model that can be fine-tuned on downstream tasks. To further improve performance, other models incorporate structural information [19, 20], exploit the GO hierarchy [9, 21], or ensemble deep-learning outputs with traditional similarity-based approaches [9, 16].

Both homology-based and existing *de novo* methods, however, suffer from meaningful drawbacks. First, both classes of methods can only predict the static set of functions in their database training set. With an average of 298 GO terms introduced and 591 GO terms deprecated each year (Fig. S1), models become outdated almost immediately. Moreover, both approaches overlook the textual descriptions accompanying the labels, a rich source of information that may improve performance, especially on rare functions. While models like ProteinCLIP [22] leverage sequence text descriptions and contrastive learning to enhance applications such as prediction of protein-protein interactions and homology detection, they are not yet tested for any type of function prediction. To bridge these gaps, recent methods for few- and zero-shot function prediction have been proposed [23, 24]. Few-shot function prediction consists of predicting functions that are represented only by a small number of sequences in the training set, whereas zero-shot prediction aims to predict functions that are not present in the training set, including new functions that no protein has been annotated with. These methods typically enable zero-shot prediction by incorporating additional information during inference such as label ontology relationships [24], gene network topology [23], and textual descriptions [23]. While these approaches improve performance on low-frequency labels, they underperform or are untested on traditional prediction with known labels, rely on auxiliary data that may not be available during inference, and are evaluated on artificial zero-shot test sets that overlap in time with the training data. To better simulate inference scenarios and avoid data leakage, an ideal zero-shot test set would consist of data collected after the training set date cutoff. Such an approach more accurately reflects how protein function prediction models are used in practice and mirrors best-practices from other studies such as the CAFA challenge [25].

We hypothesized that leveraging multimodal inputs – by extracting knowledge from both a protein’s amino acid sequence and the text description of its function – could enable strong performance on protein function prediction in both the supervised and zero-shot settings. We couple semantic protein embeddings with text embeddings from a Large Language Model (LLM) to train ProtNote, the first deep learning model capable of both supervised and zero-shot protein function prediction by training over both amino acid sequences and free-form text information (Fig. 1B-D). ProtNote matches the performance of ProteInfer, the current state-of-the-art GO annotation prediction model, in the supervised setting and crucially outperforms baselines in the zero-shot setting. We demonstrate how ProtNote captures nuanced sequence-function relationships and unlocks a range of biological use cases inaccessible to prior models.

## Materials and Methods

### Predicting protein-text likelihood

To build a model capable of both supervised and zero-shot protein-function annotation, we formulate GO term prediction from protein sequence as a binary classification problem: given a protein-GO term pair, the model predicts if the protein has the presented GO term. Class imbalance, however, posed a challenge, as 80% of the labels are used only in 0.03% of the training sequences (Fig. S3). We employ a two-fold approach to address this obstacle.

First, to accelerate learning from hard examples, we apply a focal loss with *γ* = 2 and no *α* parameter. Let *λ*_*j*_ denote GO term *j* and *x*_*i*_ represent protein sequence *i*, where *j* ∈ {1, 2, …, *L*}, *i* ∈ {1, 2, …, *N* }, with *L* and *N* being the total number of GO terms and sequences, respectively. Let *p*_*i,j*_ ∈ [0, 1] denote the model’s estimated probability that sequence *x*_*i*_ is annotated with go term *λ*_*j*_. The focal loss is defined as:

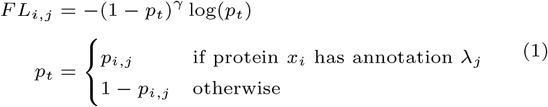

Second, we sample protein sequences during training according to the inverse frequency of their functional annotations. Let *f*_*j*_ be the frequency of GO term *λ*_*j*_ in the training set, then the sampling weight *w*_*i*_ of a sequence *i* is given by:

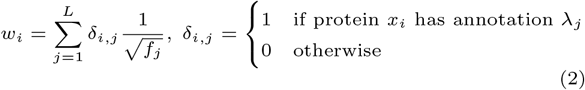

We use the square root of the frequency to avoid unreasonably high weights resulting from the extreme label imbalance.

### Datasets

ProtNote is trained on descriptions from GO annotations. These annotations capture our knowledge of three aspects of protein biology: molecular function, biological process, and cellular component. Consequently, GO terms are phrases describing the molecular actions of gene products, the biological processes in which those actions occur, and the cellular locations where they are present. We train our model with the GO terms descriptions from the 01 July 2019 GO Annotations release.

Within the supervised setting, we leverage ProteInfer’s [9] random split, where 80%, 10%, and 10% of sequences were assigned to train, validation, and test, respectively. This dataset is constructed from the SwissProt [26] section of the UniProt [5] database, the world’s largest repository of manually curated, high-quality protein sequence and function information. We remove duplicated sequences and long sequences with more than 10,000 amino acids.

Since our method extends ProteInfer’s original weights and architecture, we refer to the GO terms it was trained on as *in-vocabulary* and all others as *out-of-vocabulary*. GO terms live in a directed acyclic graph, where any term can have multiple ancestors. However, UniProt entries only show the most specific known GO term, ignoring all ancestor terms. Using only the most specific terms during training would teach the model to predict positive for the specific label but negative for the parents. Therefore, we follow ProteInfer’s annotation scheme, where *all* terms – both the most specific and the ancestors – associated with a sequence are used during training.

To evaluate zero-shot performance on unseen, new GO annotations, we create a GO zero-shot test set from the 17 June 2024 GO Annotations release and the May 2024 SwissProt [26] release; this zero-shot set includes (1) only sequences that were added to SwissProt between July 2019 (the version used by ProteInfer) and May 2024 and (2) only out-of-vocabulary GO terms, i.e., functions that were first described within the GO ontology after July 01, 2019. Figure S2 quantifies the effect of changes in the GO graph on model performance.

To explore ProtNote’s zero-shot capabilities on an out-of-distribution task, we test its performance on prediction of EC numbers. We directly use ProteInfer’s EC numbers test set, consisting of sequences and annotations that ProtNote was not trained on, and extract the functional text descriptions from ExPASy [27]. EC numbers are composed of four levels: the first level indicates the enzyme class based on the type of reaction catalyzed; the second and third levels correspond to the enzyme subclass and sub-subclass, respectively, providing more detail on the type of molecular group, bond, or product involved; and the fourth level is the enzyme identification, representing the specific enzyme and its relationship to particular metabolites or cofactors. We create a single textual description for each EC number by concatenating the descriptions of all its levels, separated by commas. Statistics for all datasets and splits are specified in Table 1.

**Table 1.** Dataset Statistics. GO-Rand-Train, GO-Rand-Val, and GO-Rand-Test splits are based on those from ProteInfer [9]. Not all possible GO terms are observed in all splits, and there are about 16% duplicated sequences. The length of sequences and number of annotations have extremely skewed distributions. GO-Zero-Shot and GO-Zero-Shot-LN correspond to the datasets with unseen GO annotations using all the labels and only the leaf nodes, respectively. The GO Zero Shot datasets have significantly fewer labels per sequence, as they only include the new labels introduced in the 2024 GO release. EC-Zero-Shot consists of unseen sequences with EC annotations, which ProtNote was not trained on. See Methods for additional details.

| Split | Seqs | Unique Seqs | Median Seq Length | Max Seq Length | Median $\frac{\text{Labels}}{\text{Seq}}$ | Max $\frac{\text{Labels}}{\text{Seq}}$ | Unique Labels |
| --- | --- | --- | --- | --- | --- | --- | --- |
| GO-Rand-Train | 418015 | 351429 | 303 | 35213 | 43 | 1008 | 31365 |
| GO-Rand-Val | 52841 | 44362 | 303 | 10061 | 43 | 760 | 21914 |
| GO-Rand-Test | 51751 | 43575 | 301 | 11103 | 43 | 815 | 22026 |
| GO-Zero-Shot | 63594 | 56212 | 303 | 15639 | 1 | 14 | 614 |
| GO-Zero-Shot-LN | 2445 | 2276 | 430 | 7158 | 1 | 6 | 402 |
| EC-Zero-Shot | 25892 | 21870 | 347 | 8903 | 4 | 22 | 2534 |

### ProtNote architecture, training, and inference

An overview of the ProtNote architecture is presented in Figure S4. ProtNote uses ProteInfer [9] as a protein sequence encoder and the Multilingual E5 Text Embedding model (E5) [28] as a text encoder – both encoders are frozen throughout training. To transform the embeddings to a more suitable space and reshape them to a common size *d* = 1024, they are each passed through independent multi-layer perceptrons (MLP) with three hidden layers of 3*d* = 3*x*1024 = 3072 units each and an output layer of size *d* = 1024. These embeddings are concatenated and fed through an MLP with three hidden layers of 3*d* = 3072 units each, followed by the output neuron. The MLPs employ ReLU activations [29] and have no bias parameters because batch normalization [30] is used.

While we employ E5, there are several options for the text encoder, and ProtNote’s architecture is flexible to the choice of text (and protein sequence) encoder. We chose E5 Text embeddings over the popular biomedical-specific model BioGPT [31], since we empirically observed improved performance with E5 as a label encoder (Fig. S5). We speculate that this could be due to E5’s high-quality sentence embeddings [28]; to the best of our knowledge, there are no biomedical-specific models that are explicitly designed to produce sentence embeddings – models like BioGPT extract sentence embeddings by averaging token-level embeddings.

During training, ProtNote uses a key set of strategies to regularize learning, generalize to rare annotations, and avoid overfitting. First, we corrupt the protein sequences by randomly applying conservative residue substitutions using the BLOSUM62 [32] matrix. Second, we employ focal loss [33] and weighted sampling to mitigate the extreme label imbalance in GO annotations. There are over 32,000 annotated functions in GO, most of them with extremely low frequency; thus, without focal loss and sequence weighted sampling, the model’s natural tendency would be to focus on the most frequent classes and predict the rest of the labels as negatives. Third, the Gene Ontology offers both short and long descriptions for each GO term; we utilize both descriptions as a form of data augmentation. During training, each time an annotation is encountered, we randomly sample one of these descriptions. In contrast, during inference, we run ProtNote twice for a given annotation – once each for each of the short and long descriptions – and ensemble the probabilities by averaging them. Sometimes, GO terms can have more than two descriptions called “synonyms”; because not all terms have synonyms, for simplicity we do not use them. For EC numbers, only one description is available per annotation, so the ensemble procedure simplifies to running ProtNote once per annotation. Finally, we inject random noise to the label encoder embeddings before the initial MLP. More details about these techniques are available in the Supplementary Information.

ProtNote is trained for 46 epochs on 8x 32GB V100 NVIDIA GPUs using an effective batch size of 256 (32 × 8) with dynamic padding. The model trains using the Adam optimizer [34] with a learning rate of 0.0003, employs gradient clipping set to 1, and uses mixed precision [35]. We select the model checkpoint based on best validation performance.

### Baselines

In the supervised setting, we directly compare ProtNote against ProteInfer and BLAST and use ProteInfer’s test set. The BLAST baseline identifies the most similar sequence in the training set (i.e., the top BLAST hit) for each query sequence in the test set and then uses the labels of the top hit as the predicted labels for the query sequence.

To our knowledge, no prior model has evaluated zero-shot prediction of protein function from textual label descriptions alone. Thus, in the zero-shot setting, we devised a novel benchmark, akin to a “label description similarity baseline”, to lower bound our model’s performance. First, we map each out-of-vocabulary term to the closest in-vocabulary term based on the function description’s cosine similarity using E5 or BioGPT text embeddings. Then, we directly use ProteInfer to predict likelihoods for the in-vocabulary labels and simply propagate these predictions to the new labels based on the created mapping.

### Evaluation metrics

To evaluate the performance of different models, we use both macro- and micro-averaged mean Average Precision (mAP), also known as the Area Under the Precision-Recall Curve (AUPRC). The mAP metric summarizes model performance across all possible prediction thresholds, eliminating threshold tuning and providing a more robust assessment of model performance than threshold-dependent metrics such as *F*_1_ or *F*_*max*_ scores [25]. We report both mAP Macro and mAP Micro because of their different virtues, although we use mAP Macro for model selection. Macro averaging offers a performance estimate that is unbiased by the GO term distribution, ensuring that each label contributes equally to the final score regardless of its prevalence in the dataset. This is particularly important given that the distribution of GO terms across sequences is extremely skewed, with 98% of GO terms appearing for at most 1% of the sequences, while the 10 most frequent labels are present for at least 58% of the sequences (Fig. S3). On the other hand, micro averaging is influenced by the label distribution, which can be important when the goal is to optimize for the most frequently occurring outcomes.

## Results

ProtNote is a deep-learning model capable of both supervised and zero-shot protein function prediction using only protein amino acid sequences and text-based functional descriptions as input. Our primary aim was to design a model capable of zero-shot inference – that is, predicting new functions that lie outside of the training set and, ultimately, that no protein has been annotated with. To achieve this, we train ProtNote to predict the likelihood that a protein is annotated with a specific function given its amino acid sequence and functional text description (Fig. 1B). We leverage ProteInfer [9] and E5 Text Embeddings [28] to separately encode the sequence and text, respectively, and then fuse these embeddings for prediction.

We evaluate ProtNote in both the supervised (Fig. 1C) and zero-shot (Fig. 1D) settings, benchmarking against the state-of-the-art deep-learning method ProteInfer and the gold-standard, homology-based method BLAST in the supervised setting and against custom embedding-based baselines in the zero-shot setting. For completeness, we break down the models’ performance across the three GO Ontologies (biological process, cellular component, molecular function) (Figs. S6, S7) and the seven top-level EC classes (oxidoreductases, transferases, hydrolases, lyases, isomerases, ligases, translocases) (Fig. S8).

### ProtNote meets state-of-the-art performance in supervised GO function prediction

To evaluate the performance of ProtNote, we first tested the model in the supervised setting, using mean Average Precision (mAP) metrics to compare our method to ProteInfer [9], the current leading model for GO annotation prediction (Fig. 2A). Overall, ProtNote matches ProteInfer’s and exceeds BLAST’s performance on the test set (Fig. 2B-C) despite using a more general and scalable (Fig. S9) architecture capable of zero-shot inference. ProteInfer retains a minor mAP Macro advantage over our model (Fig. 2B; avg. mAP Macro of 0.6417 vs. 0.6018 for ProteInfer vs. ProtNote, respectively; two-sided Welch’s t-test *p <* 0.0001), while ProtNote is on par for mAP Micro (Fig. 2C; avg. mAP Micro of 0.9032 vs. 0.9042 for ProteInfer vs. ProtNote, respectively; two-sided Welch’s t-test *p* = 0.2298). These mAP Micro and Macro observations are consistent across the three GO Ontologies (Fig. S6), except for the mAP Macro results for the molecular function ontology, where BLAST has a mild advantage over ProtNote (0.7292 vs. 0.7080 for BLAST vs. ProtNote, respectively). Notably, during our experiments, we observed a trade-off between supervised and zero-shot performance (Fig. S5). Eliminating label embedding noise and description sampling increased ProtNote’s performance in the supervised setting, even surpassing ProteInfer on both mAP Micro and mAP Macro metrics, but at the expense of zero-shot capabilities (Fig. S5).

**Fig. 2.**
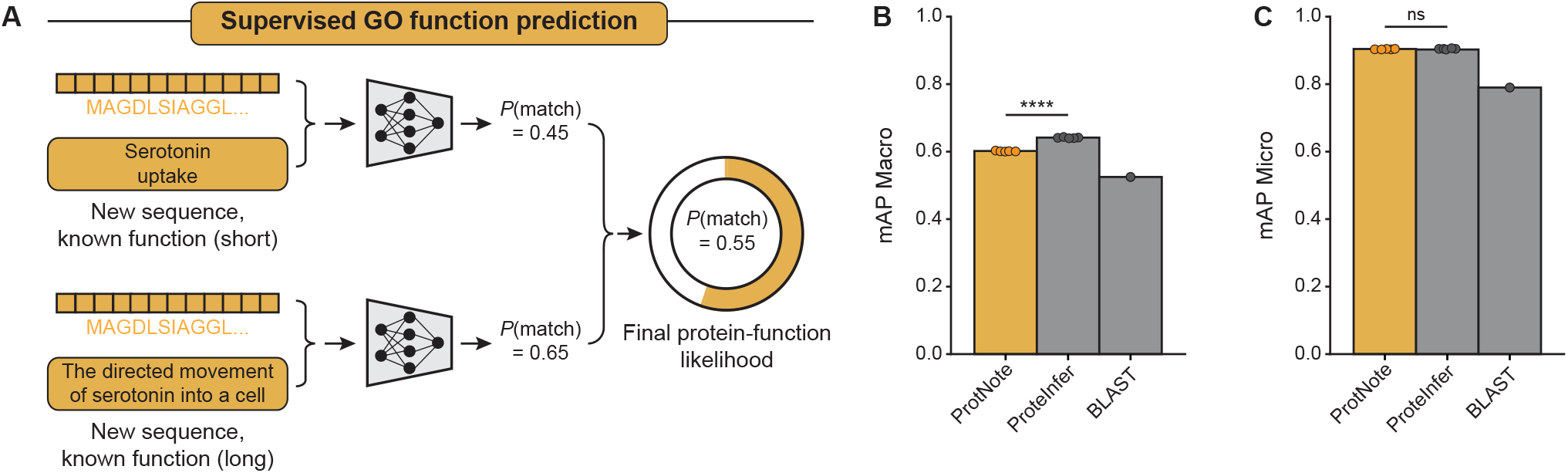
ProtNote achieves robust supervised prediction of GO function annotations. **(A)** In the supervised setting, ProtNote predicts protein-function likelihoods for unseen sequences with known GO annotations. ProtNote is used twice for each protein-function pair, once for the function’s short textual description and another for the long textual description. The final likelihood score is the average of the scores obtained from both descriptions. **(B-C)** mAP Macro (B) and mAP Micro (C) scores for GO annotation prediction in the supervised setting, comparing ProtNote (yellow; *n*=5 independently trained models) against ProteInfer (grey; *n*=5 seeds) and BLAST (grey) (mean *±* s.d.; two-sided Welch’s t test, *p*^∗∗∗∗^ *<* 0.0001, *p*^ns^ = 0.230).

### ProtNote’s embedding space captures protein functional relationships

Motivated by the model’s performance in the supervised setting, we next inspected ProtNote’s multimodal embedding space by reducing the dimensionality using uniform manifold approximation and projection (UMAP) (Fig. 3). We observe that positive protein-function pairs (i.e., human-verified SwissProt GO annotations) group together in a small region of the embedding space (Fig. 3A), suggesting that ProtNote learns to group semantically similar pairs closely and identifies true underlying functional relationships in the protein data. Looking at the joint embeddings from only positive protein-function pairs (Fig. 3B), we observe that embeddings cluster by GO ontology groups, indicating that ProtNote’s embeddings additionally capture more nuanced biological information on function. It is worth noting that ProtNote is only trained to predict whether a pair matches and is not explicitly trained (e.g., in a contrastive manner) to differentiate among the different GO ontologies.

**Fig. 3.**
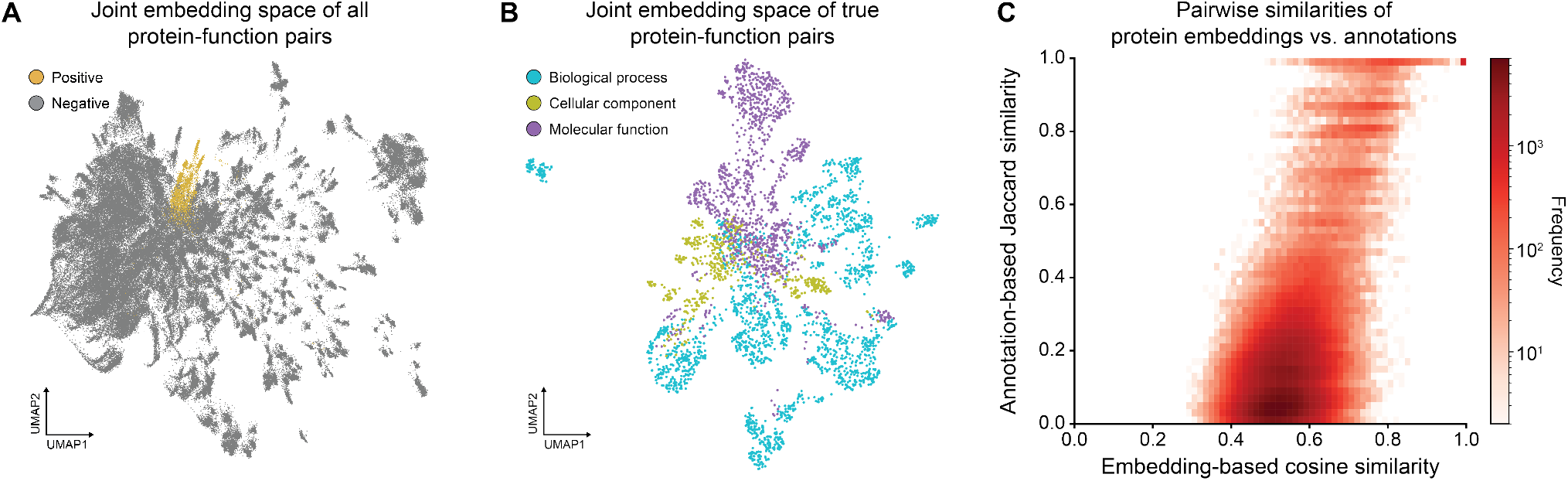
ProtNote embeddings capture functional relationships. All plots are based on random samples of the test set consisting of the most frequent GO annotations (*n*=1,731). **(A)** UMAP projection of ProtNote output layer embeddings for protein-function pairs, where color indicates if the sequence is paired with a true positive (yellow) SwissProt GO annotation or negative (grey) label (*n*=100 proteins x 1,731 annotations = 1,731,000 pairs). **(B)** UMAP projection of ProtNote embeddings for the positive protein-function pairs from (A), where color indicates the GO Ontology of the pair’s function (*n*=30,436). **(C)** 2D histogram of the pairwise cosine similarity between amino acid sequence embeddings and the pairwise Jaccard similarity between sequence annotations for the protein-protein pairs of the sampled test set sequences (*n*=800 proteins x 800 proteins = 640,000 pairs).

Based on these observations, we hypothesized that similar protein sequences would tend to have similar functional labels, and that ProtNote protein embeddings from the sequence projection head would reflect this relationship. To investigate this, we first define two ways to capture the similarity between a pair of proteins. First, we define the embedding-based similarity as the cosine similarity between the output embeddings from the amino-acid sequence projection head. This similarity metric carries no explicit information about functional annotations. Second, we specify the annotation-based similarity as the Jaccard similarity coefficient of the two proteins’ multi-hot annotation vectors. Thus, the annotation-based similarity captures no explicit sequence information. We observe a correlation between the embedding-based and annotation-based pairwise similarities (Fig. 3C; Pearson *R*=0.533, *p <* 0.0001), implying that even before the output MLP, sequence embeddings from the projection head already contain significant functional information. We also observe a mild correlation between pairwise sequence and label embedding similarities (Fig. S10; Pearson *R*=0.404, *p <* 0.0001), revealing that higher sequence similarity is associated with higher functional label similarity, even though the model does not explicitly align the sequence and text embeddings.

### ProtNote enables zero-shot function prediction

ProtNote is designed to enable zero-shot inference when given a set of unseen protein sequences and novel functional descriptions. We thus tested ProtNote in two zero-shot tasks: predicting protein functions based on novel GO annotations and predicting EC numbers.

#### Zero-shot prediction of novel GO annotations

To assess ProtNote’s ability to predict novel GO annotations for unseen protein sequences (Fig. 4A), we evaluated its performance in two settings: on GO annotation leaf nodes only, and on both leaf nodes *and* inferred nodes (Fig. 4B-C, Fig. S7). Leaf nodes refer to GO terms with no children, while inferred nodes designate terms that can be derived from another annotation based on the graph hierarchy. For example, the GO terms “biological regulation” (GO:0008150) and “biological process” (GO:0065007) can be inferred from the term “regulation of biological process” (GO:0050789), as the first two are parents of the latter. In general, when measuring zero-shot capabilities, we consider only the net-new inferred nodes.

**Fig. 4.**
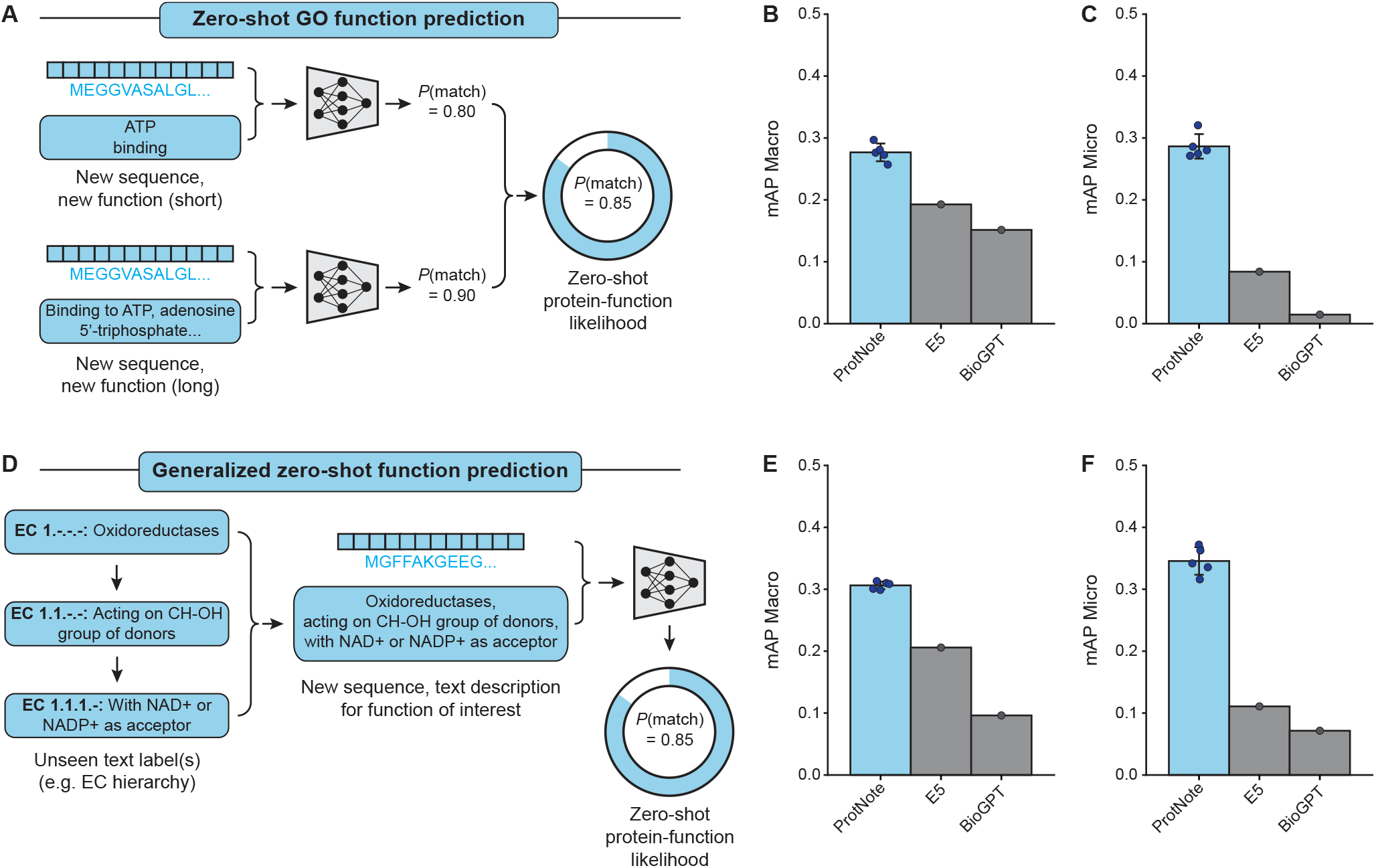
ProtNote enables generalizable zero-shot protein function prediction. **(A)** In zero-shot GO function prediction, ProtNote predicts protein-function likelihoods for unseen sequences with new, unseen GO annotations. As in the supervised setting, ProtNote is used twice for each protein-function pair, and the final likelihood score is the average of the scores obtained from both short and long textual descriptions. **(B-C)** mAP Macro (B) and mAP Micro (C) scores for zero-shot prediction of novel GO annotations for the GO leaf nodes, comparing ProtNote (blue; *n*=5 independently trained models; mean *±* s.d.) against the Multilingual Instruct E5 and BioGPT (grey) baselines. **(D)** Demonstration of generalized zero-shot function prediction with Enzyme Commission (EC) numbers. A full textual description for each EC number is created by concatenating its description with those of its parents, separated by commas. A protein sequence and its corresponding aggregate text description are input into ProtNote to predict a protein-function likelihood score. **(E-F)** mAP Macro (E) and mAP Micro (F) scores for zero-shot prediction of EC numbers, comparing ProtNote (blue; *n*=5 independently trained models; mean *±* s.d.) against the E5 and BioGPT (grey) baselines.

ProtNote outperforms baselines across all scenarios evaluated (Fig. 4B-C, avg. mAP Macro of 0.2139, 0.1556, 0.1236 for ProtNote, E5, BioGPT respectively; Fig. S7). While ProtNote exceeds the baselines on all metrics in the scenario with just leaf nodes (Fig. 4B-C), we observe the largest performance gap when comparing ProtNote’s predictions with baseline mAP Micro metrics in the scenario including both leaf and inferred nodes (Fig. S7D), suggesting that the baselines perform poorly on the most frequent annotations. We note that a large portion of the performance drop between the supervised and zero-shot settings may be explained by the change in the inferred labels caused by changes in the GO graph (Fig. S2).

We also observe significant variation in performance by GO ontology (Fig. S7). For example, most models excel at predicting labels that fall within the “molecular function” ontology and under-perform when predicting sequence annotations from the “cellular component” ontology. We hypothesize that this discrepancy could be understood from two angles: the annotations and the sequences. First, textual annotations bias towards describing molecular functions and biological processes rather than cellular components. Only 14% of the positive GO terms in the training set come from the cellular component ontology, making it the least represented ontology. Second, prior work has suggested that protein language models may learn evolutionarily-conserved motifs through self-supervised pretraining [36], but must be fine-tuned for prediction of subcellular location specifications [37] that may be determined by a small number of characteristic residues.

#### Zero-shot prediction of EC numbers

To further assess ProtNote’s zero-shot capabilities in a markedly different task, we applied it to predicting EC numbers – a task the model was not trained on. This task involves predicting labels that describe enzyme functions using a distinct terminology that does not directly map to GO annotations (Fig. 4D). For each EC number, we constructed the input text by concatenating its description with those of its parents, separated by commas (Fig. 4D). We then used only this final text for inference without ensembling. Finally, we assessed the performance of our model across the top-level EC classes and compared it against the E5 and BioGPT baselines (Fig. 4E-F, Fig. S8).

ProtNote outperforms the E5 and BioGPT models, both for mAP Macro (Fig. 4E; avg. scores of 0.3062, 0.2058, 0.0961 for ProtNote, E5, BioGPT respectively) and mAP Micro (Fig. 4F; avg. scores of 0.3456, 0.1107, 0.0714 for ProtNote, E5, BioGPT respectively). Similar to the zero-shot GO annotation task, the baselines show much lower mAP Micro than mAP Macro metrics, reflecting their poor performance on frequent EC numbers. Furthermore, ProtNote demonstrates superior performance across all seven top-level EC classes (Fig. S8). Notably, all models perform well on ligases but struggle on translocases (Fig. S8).

Together, these results demonstrate that ProtNote performs robustly in zero-shot prediction of protein function annotations, as evidenced by both extrapolation to unseen GO annotations and function prediction in a completely unseen scenario of EC descriptions. While ProtNote does not reach the metrics from the fully supervised setting, as expected, its strength in zero-shot settings highlights the effectiveness of combining amino-acid sequence and text data to predict protein functions.

## Discussion

In this work, we introduce ProtNote, a multimodal deep learning model capable of supervised and zero-shot protein function annotation. We demonstrate how ProtNote leverages unstructured text to predict the likelihood that proteins perform arbitrary functions. Importantly, we showcase that ProtNote generalizes to unseen sequences and functions by evaluating the model on newly-added GO annotations and on enzyme annotations, which were not used to train the model. We observe that this generality does not come at the cost of supervised performance; our model is also performant at predicting known GO terms, performing on par with the state-of-the-art model ProteInfer.

To the best of our knowledge, ProtNote is the first deep learning model capable of both supervised and zero-shot protein function prediction using only a single sequence-text pair. Unlike homology-based methods such as BLAST [6] and profile Hidden Markov Models [7, 8], ProtNote does not rely on direct comparisons against thousands of other sequences. Instead, ProtNote depends solely on a protein’s amino acid sequence and a text description for zero-shot inference. This flexibility contrasts with recent machine-learning based attempts that depend on multiple – and often scarce – inputs from different modalities [23, 24] and that evaluate zero-shot capabilities based on synthetically crafted datasets rather than real time-based splits. Compared to traditional supervised approaches, ProtNote is more robust to changes in the GO, as it is less impacted by new, removed, or redefined terms.

Despite its demonstrated performance and versatility, ProtNote leaves significant possibilities for future improvement and expansion. We train only on the GO dataset, which inherently introduces several obstacles. First, while ProtNote is more robust to changes in the GO relative to supervised approaches trained on a fixed set of terms, like all GO-trained models, it remains prone to prediction errors driven by changes in the GO graph. As shown in Fig. S1, changes in the graph structure directly modify the annotations implied by GO ancestry. Second, the model might overfit to text descriptions with similar content and style to those observed in the Ontology. The sequence-function matching formulation also has some drawbacks. To make predictions, ProtNote must run inference through an MLP for every protein-text pair, requiring more computational resources than other strategies. Further, ProtNote does not leverage relationships between functions to improve its predictions.

There are a number of promising research angles that could be explored to improve model performance and enable additional applications. First, training text may be expanded from GO function annotations to include other categories such as subcellular location, catalytic activity, involvement in diseases, and membership in protein families [5, 13, 38, 39, 40, 17]. Second, ProtNote’s joint MLP could be removed in favor of using a contrastive learning approach with LLM fine-tuning for faster inference, while learning a single representation in a joint embedding space. Third, a more principled weighted sampling scheme could be used instead of weighting by the sum of individual GO term weights. Finally, the development and use of different text encoders could be explored. We tested general-domain sentence transformers such as E5 [28] and biomedical-specific masked language models that are not trained to produce sentence embeddings [41, 31], and observed that general-domain sentence transformers yielded improved performance. Future work to train and apply biomedical-specific sentence transformers, which are currently rare, will help delineate whether biomedical-specific training provides benefits relative to general language models.

In sum, ProtNote provides a multimodal framework for protein function prediction and generalizes to functions beyond those on which it was trained. Our work is an important step towards general-purpose function prediction models that are robust to the dynamic, ever-evolving landscape of proteins.

## Data and code availability

Code is available at https://github.com/microsoft/protnote. Model weights, datasets, and computed metrics are available at https://zenodo.org/records/13897920.

## Author contributions

Conceptualization: S.C., N.C., S.A., K.K.Y., A.P.A.; Methodology: S.C., N.C., S.A., K.K.Y., A.P.A.; Software Programming: S.C., N.C.; Validation: S.C., N.C., S.A., K.K.Y., A.P.A.; Formal analysis: S.C., N.C.; Resources Provision: K.K.Y., A.P.A.; Data Curation: S.C., N.C.; Visualization: S.C., A.P.A.; Writing - Original Draft: S.C., N.C., K.K.Y., A.P.A.; Writing - Review & Editing: S.C., N.C., S.A., K.K.Y., A.P.A.; Supervision: K.K.Y., A.P.A.

## Acknowledgments

The authors thank Nicolo Fusi for helpful discussions and valuable feedback, and thank Sean Whitzell, Neil Tenenholtz, Rich Ciapala, and Hannes Schultz for assistance on using Microsoft’s compute resources.

## SUPPLEMENTARY INFORMATION

### A. Variability in the Gene Ontology (GO) and its impact on model performance

The GO is far from stable. A key aspect of the GO is that the GO Consortium is constantly refining it for the benefit of science. Despite the clear advantages of this ongoing curation, a major drawback is that the ontology changes frequently across releases, potentially affecting all downstream analyses that rely on it. This problem is of such significance that researchers have discussed it extensively and developed tools to identify and track changes in the GO [1]. There are three major sources of change in the GO that impact supervised models in different ways: GO Terms can be added or removed (Fig. S1A), their definitions may change (Fig. S1B), and their locations in the graph can evolve (Fig. S1C). Without retraining, supervised models are unable to predict new terms, while they will continue to predict old terms, introducing false positives. When a term’s definition changes, the predictions of a supervised model will have inconsistent interpretations. Furthermore, a change in the GO graph may modify the inferred labels of a given term. Although the third problem (Fig. S1C) impacts all models, the first two (Fig. S1A-B) do not alter ProtNote, since it is able to predict likelihoods for new or changed terms.

Since changes in the GO graph influence the performance of all models, we developed an evaluation to isolate and quantify this effect. We created a new test set with the same protein sequences as ProteInfer’s 2019 test set, but with updated annotations based on the May 2024 GO release. Given that the latest release contains new terms, we only kept the terms that were seen during ProteInfer’s training in its original dataset. ProteInfer’s mAP Macro and Micro decreased by 11.67 and 22.51% (absolute), respectively, on the updated test set with the same sequences but updated annotations (Fig. S2). These results demonstrate that supervised models degrade quickly for this task, highlighting the need for more flexible alternatives, and underscore the difficulty of zero-shot inference for this task.

Finally, we note that the distribution of GO terms is highly imbalanced – most of the labels are used rarely, i.e. for a small proportion of protein sequences, while the top labels in the GO are used very frequently (Fig. S3).

**Fig. S1.**
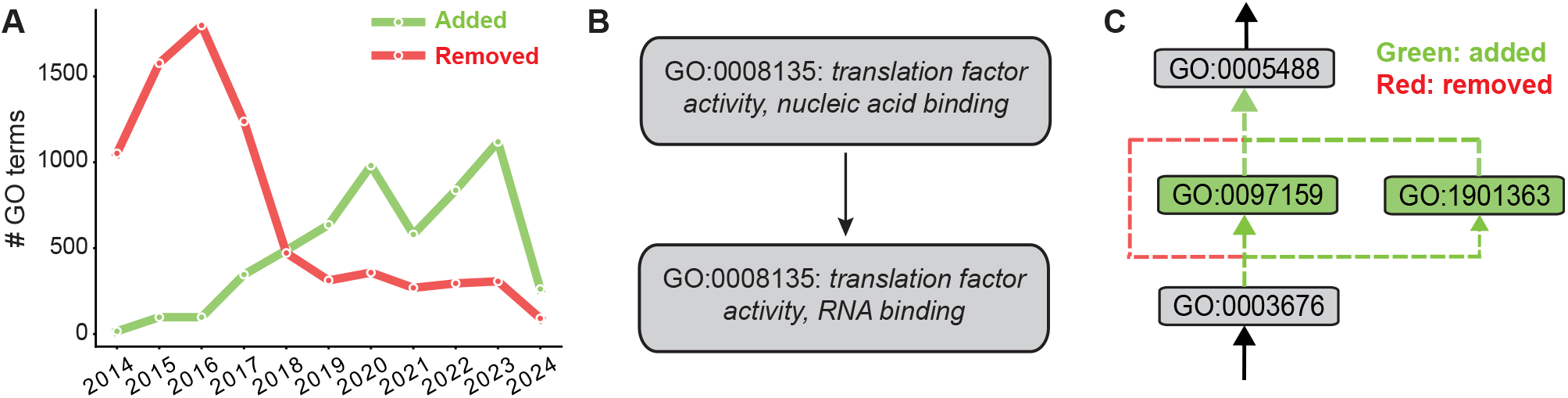
Changes in the GO are frequent and diverse. **(A)** Addition and removal of GO terms over time for the last 10 years. **(B)** Example of a GO term for which the definition changed from one release to another. **(C)** Example of a simple structure change for the parent nodes of GO term GO:0003676.

**Fig. S2.**
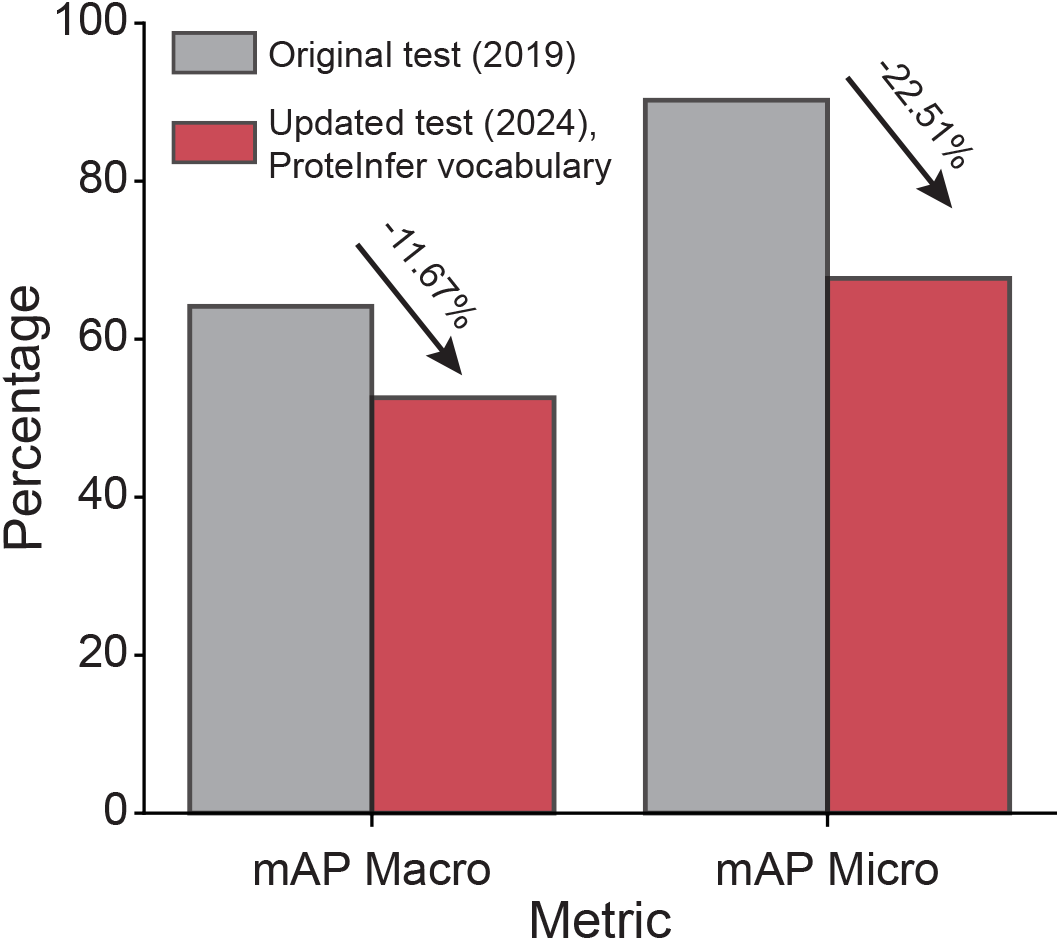
Impact of changes in the GO graph on ProteInfer baseline performance. mAP Macro and mAP Micro performance of the ProteInfer baseline on the original 2019 test set (grey) and on an updated test set (red) with the same protein sequences but with applicable GO terms based on the May 2024 GO release.

**Fig. S3.**
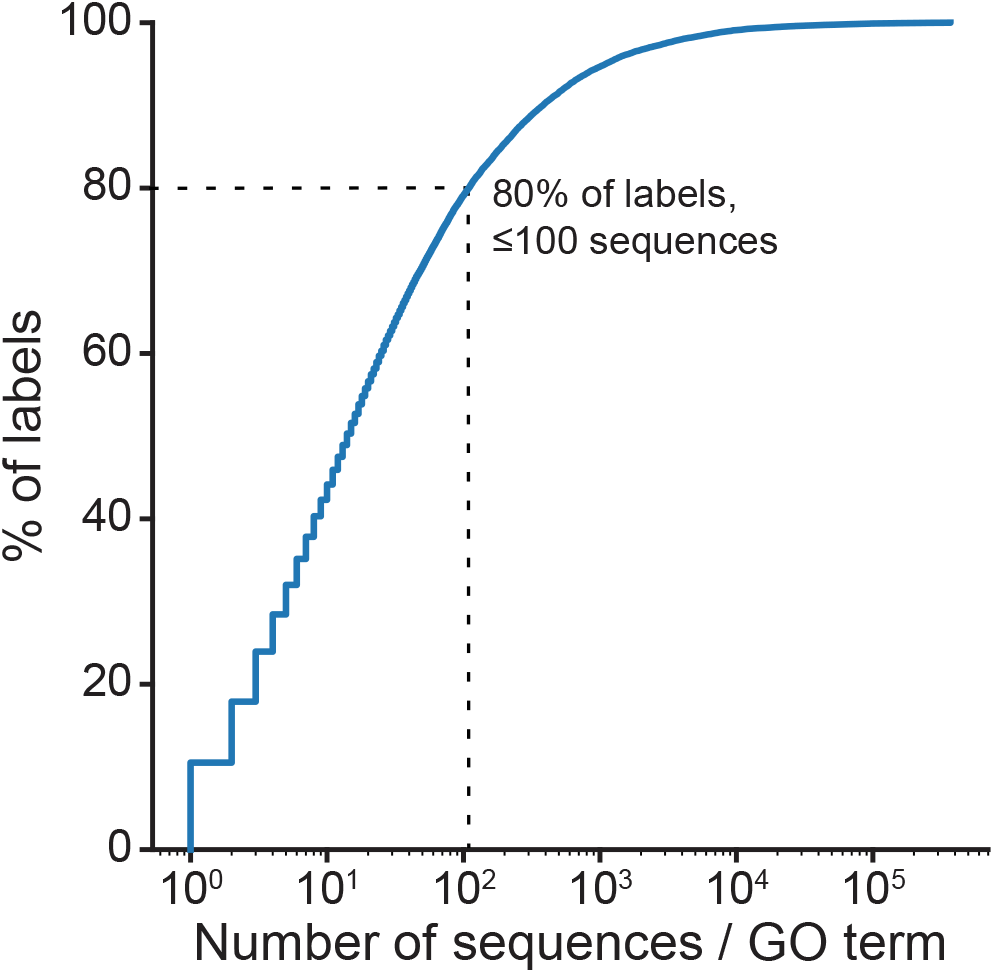
Cumulative distribution function of the number of protein sequences per GO term. The x-axis represents the number of protein sequences associated with each GO term, while the y-axis shows the cumulative proportion of GO terms used for a given number of sequences. Most of the labels are used infrequently, while the top labels are used extensively.

### B. Detailed results

A detailed schematic of the ProtNote modeling framework and architecture is provided in Figure S4. We perform a series of ablations to assess the impact of different design choices on ProtNote’s performance in both the supervised and zero-shot settings (Fig. S5). To thoroughly evaluate model performance, we assess ProtNote’s predictions across the three GO Ontologies for the supervised (Fig. S6) and zero-shot (Fig. S7) settings, and the seven top-level EC numbers (Fig. S8) in the zero-shot setting.

**Fig. S4.**
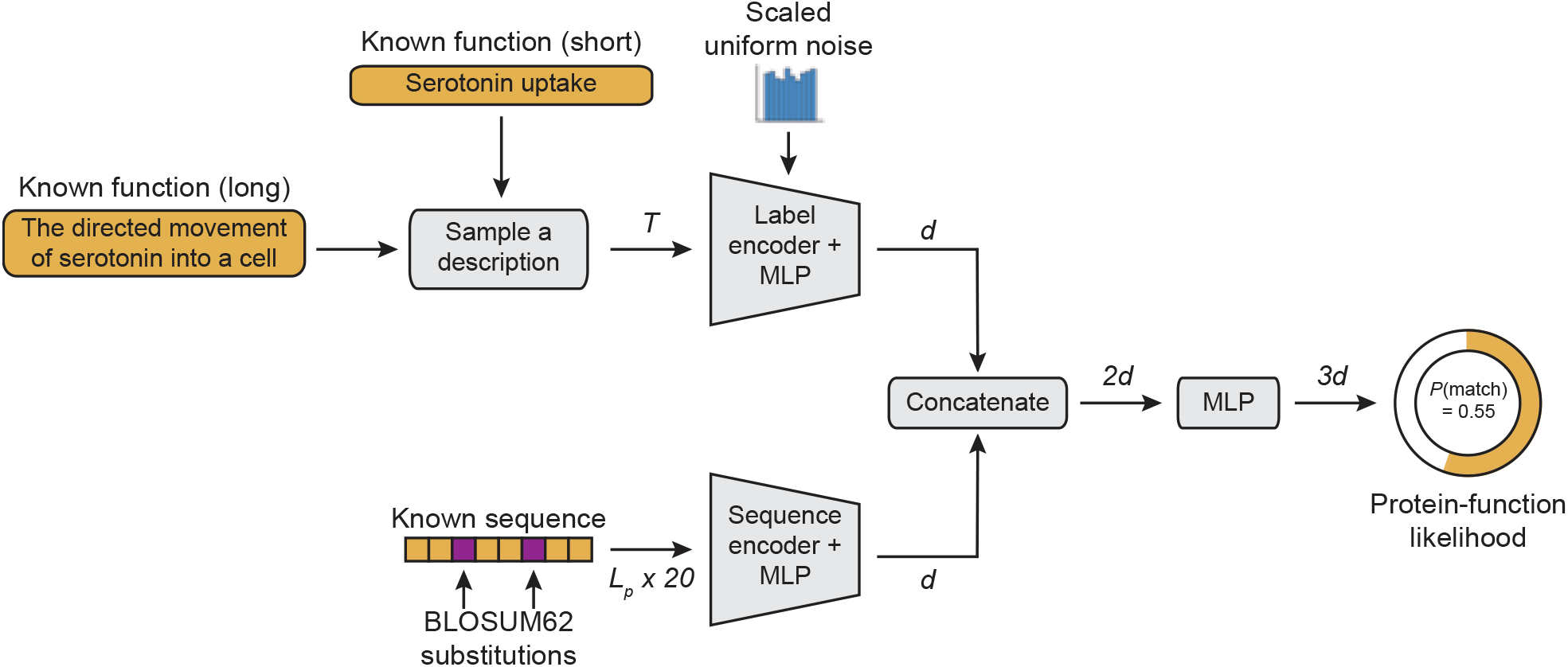
ProtNote modeling framework. During training, a protein sequence-function annotation pair is sampled. As augmentation, the protein sequence is corrupted using BLOSUM62 substitutions, and then a free-text function annotation is sampled, as either the long or short description for that function. Subsequently, the text and sequence are embedded using Multilingual E5 Text embeddings and ProteInfer, respectively, and then each embedding is projected, using a multi-layer perceptron (MLP), to a new space with common dimensionality *d* = 1024. To improve generalization noise is added to the label embeddings before the projection head. Finally, the label and sequence embeddings are concatenated and pass through a final MLP to output a protein-function likelihood.

**Fig. S5.**
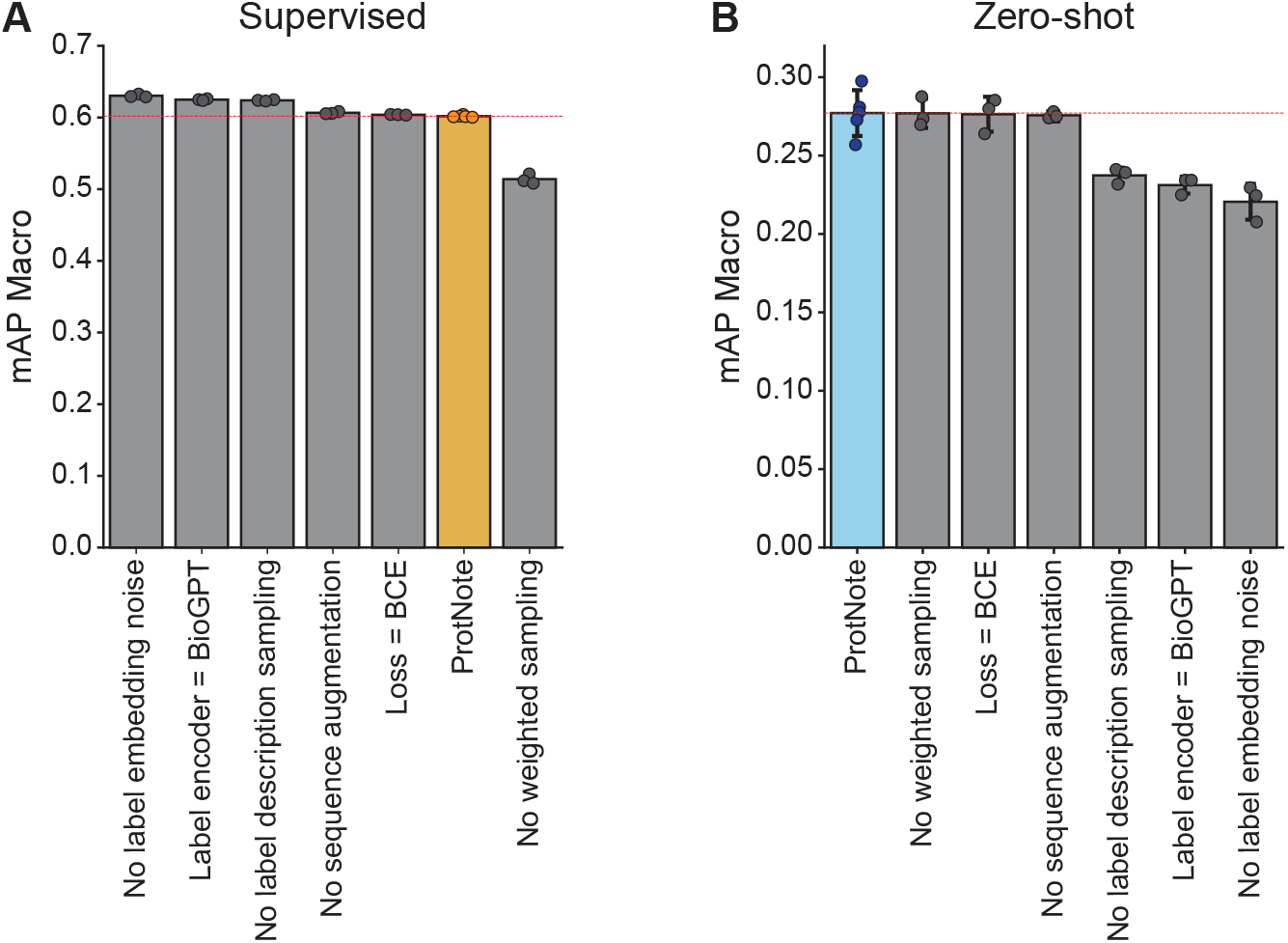
Supervised and zero-shot performance of ablated models. **(A)** mAP Macro scores for GO annotation prediction in the supervised setting, for ProtNote (yellow; *n*=5 independently trained models) and model ablations across different design choices (grey; *n*=3 independent models). **(B)** mAP Macro scores for zero-shot prediction of novel annotations for GO leaf nodes for ProtNote (yellow; *n*=5 independent models) and model ablations across different design choices (grey; *n*=3 independent models).

**Fig. S6.**
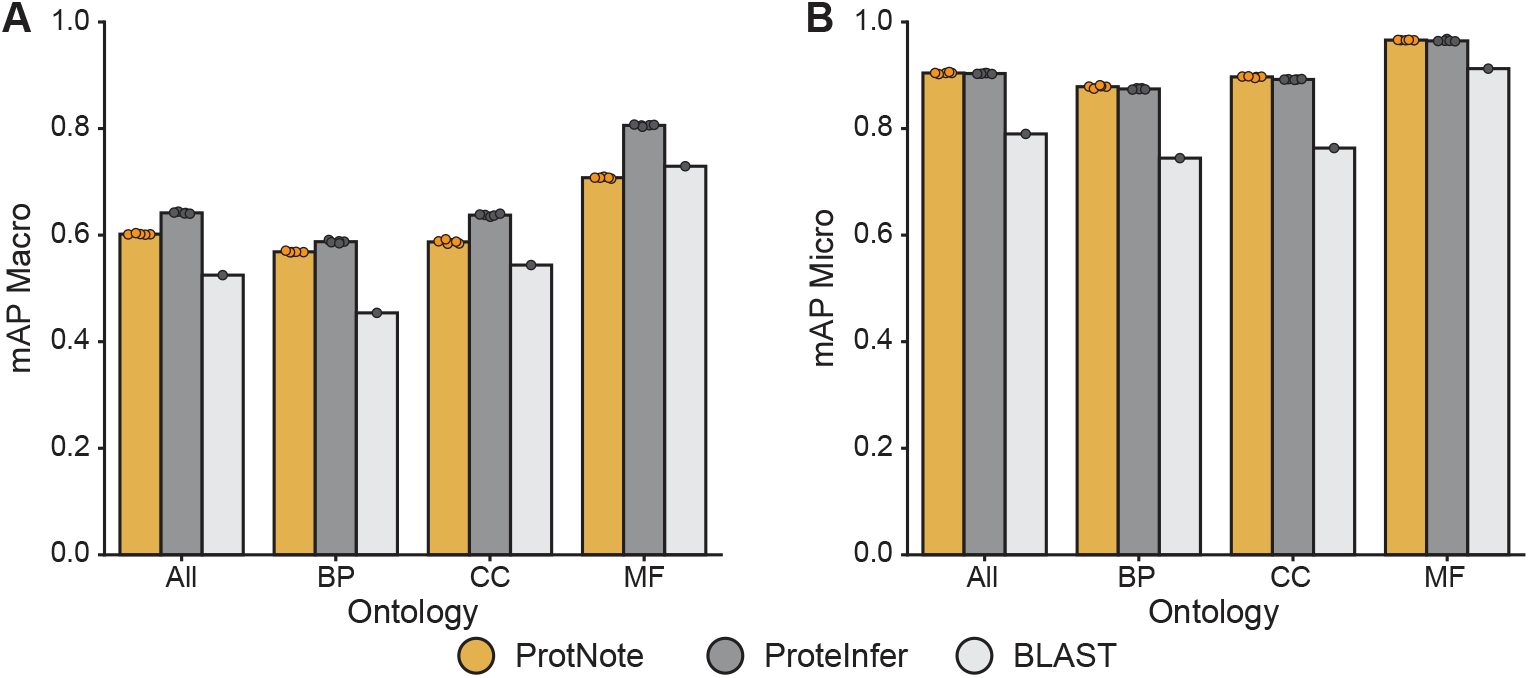
Detailed performance metrics for supervised prediction of GO function annotations over the three GO ontologies. **(A-B)** mAP Macro (A, left) and mAP Micro (B, right) scores for GO annotation prediction in the supervised setting, split by the three ontologies of the GO, Biological Process (BP), Cellular Component (CC), and Molecular Function (MF), with “All” showing the overall performance across all ontologies. ProtNote (yellow; *n*=5 independently trained models) is compared against ProteInfer (grey; *n*=5 seeds) and BLAST (white) (mean *±* s.d.).

**Fig. S7.**
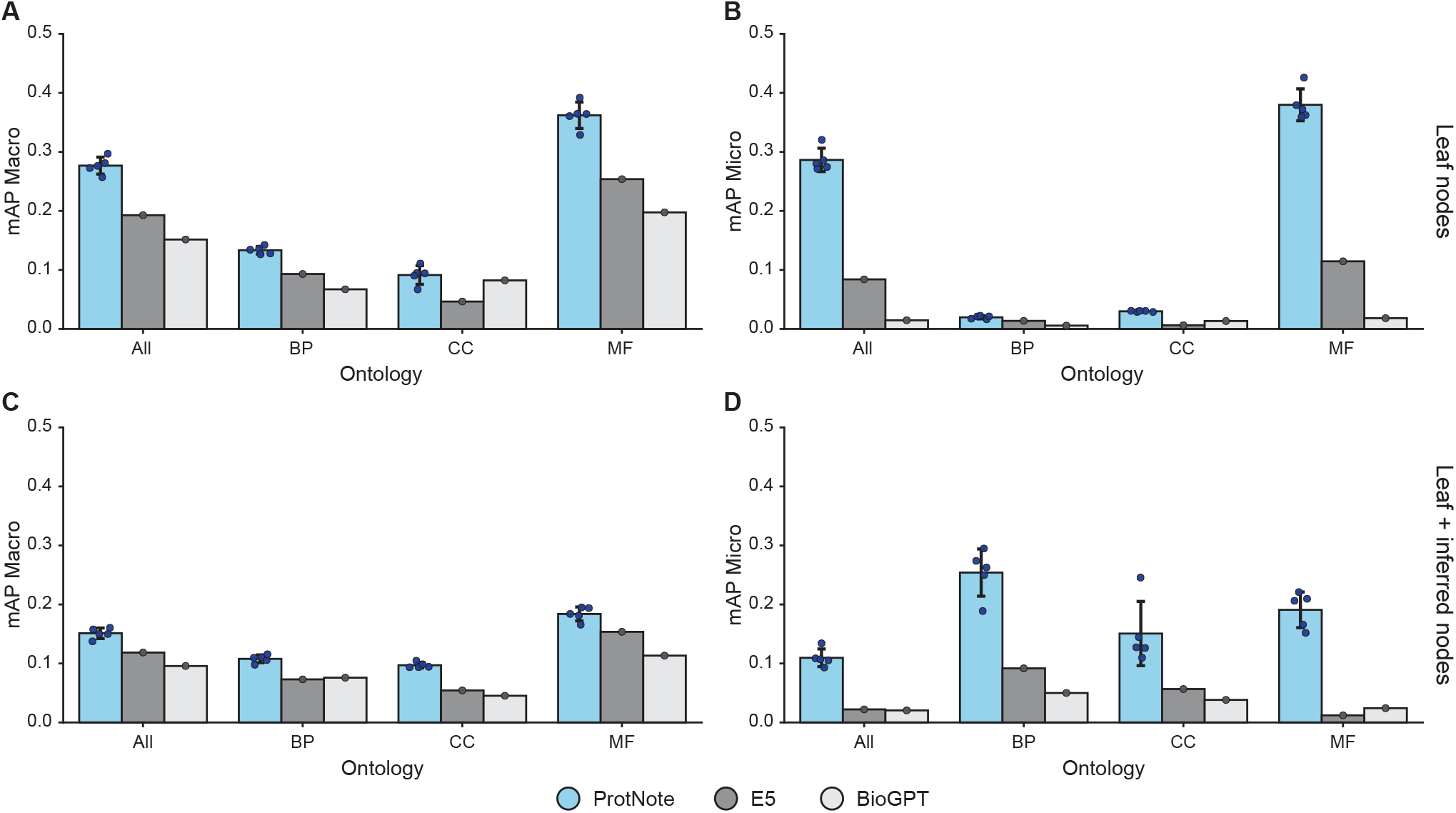
Detailed performance metrics for zero-shot prediction of novel, unseen GO function annotations over the three GO ontologies. **(A-D)** mAP Macro (left column) and mAP Micro (right column) scores for zero-shot prediction of novel GO annotations, split by the three ontologies of the GO, Biological Process (BP), Cellular Component (CC) and Molecular Function (MF), with “All” showing the overall performance across all ontologies. **(A-B)** show performance only for the GO leaf nodes; **(C-D)** show performance across all nodes: leaf nodes plus those inferred from the GO graphs. In the zero-shot setting, ProtNote (blue; *n*=5 independently trained models) is compared against the label similarity baseline using E5 (grey) and BioGPT (white) for label embedding (mean *±* s.d.).

**Fig. S8.**
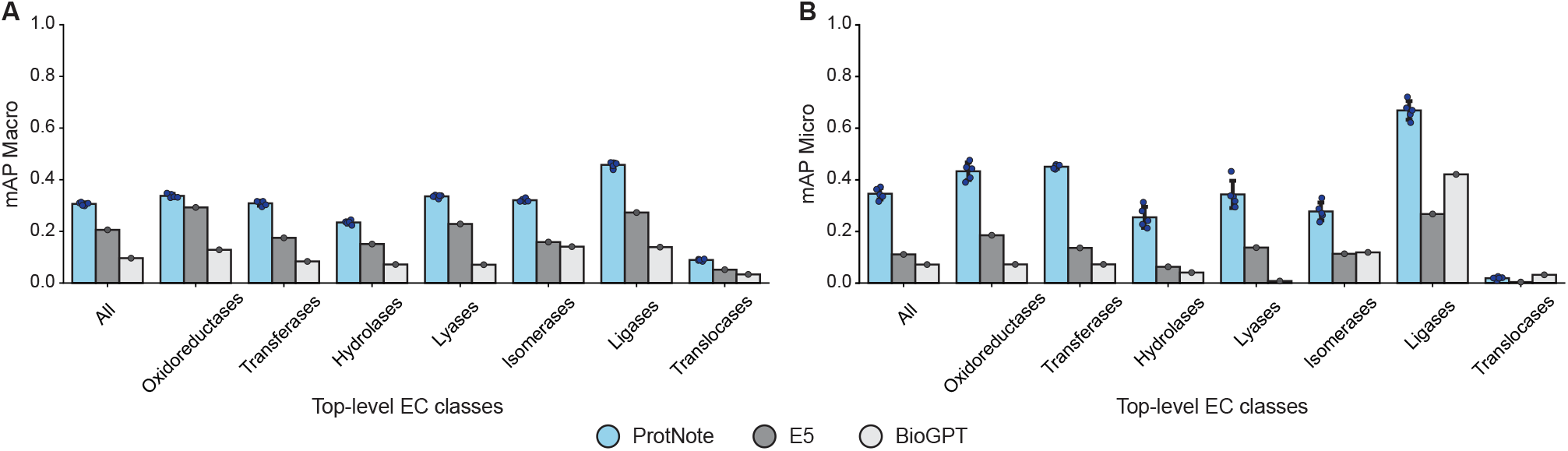
Detailed performance metrics for zero-shot prediction of Enzyme Commission (EC) numbers over the top-level enzyme classes. **(A-B)** mAP Macro (A, left) and mAP Micro (B, right) scores for zero-shot prediction of EC Numbers, split by the seven top-level EC numbers. ProtNote (blue, *n*=5 independently trained models) is compared against the label similarity baseline using E5 (grey) and BioGPT (white) for label embedding (mean *±* s.d.).

### C. Augmentation and regularization

We employ two data augmentation techniques and one regularization strategy to improve ProtNote’s generalization capacity.

#### Sequence residue substitution

To increase the robustness of ProtNote, we implement a data augmentation pipeline for all sequences, where each residue in a sequence has a probability *p* of being replaced with an amino acid sampled from the BLOSUM62 matrix, and a 1 − *p* of being left unchanged. To sample from the BLOSUM62 matrix, we select the conservative substitutions for the given amino acid, convert the substitution scores into probabilities, and then sample a substitution based on this probability distribution. The probability *p* is treated as a hyperparameter and set to 0.1 in our experiments.

#### Label description sampling

Terms in the GO database contain two attributes known as “name” and “definition”. Names are typically a one sentence description of the function, while definitions are usually a longer paragraph. Definitions are richer but often contain redundant or unnecessary information, while names have limited content but are less noisy. During our initial studies we found that using names instead of definitions resulted in better performance. However, to leverage both sources of information, we trained with both description types by randomly sampling the name or the definition of any GO term with equal probability. For inference on the validation and test sets, the model makes two predictions for each protein-function pair, one using names and the other using labels, and we ensemble these via averaging. In Fig. 2A, Fig. 4A, and Fig. S4, these are referenced as the “short” and “long” descriptions corresponding to the GO “name” and “definition” attributes, respectively.

#### Label embedding noising

Inspired by the success of introducing embedding noise during LLM fine-tuning [2] as a regularization technique, we add to the label embeddings a sample of uniform random noise in the range [− 1, 1], scaled by a factor of 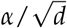. The scalar *α* is a hyperparameter and set to 20 in our experiments, and *d* is the dimensionality of the embedding space, set to 1024.

### D. Additional figures

**Fig. S9.**
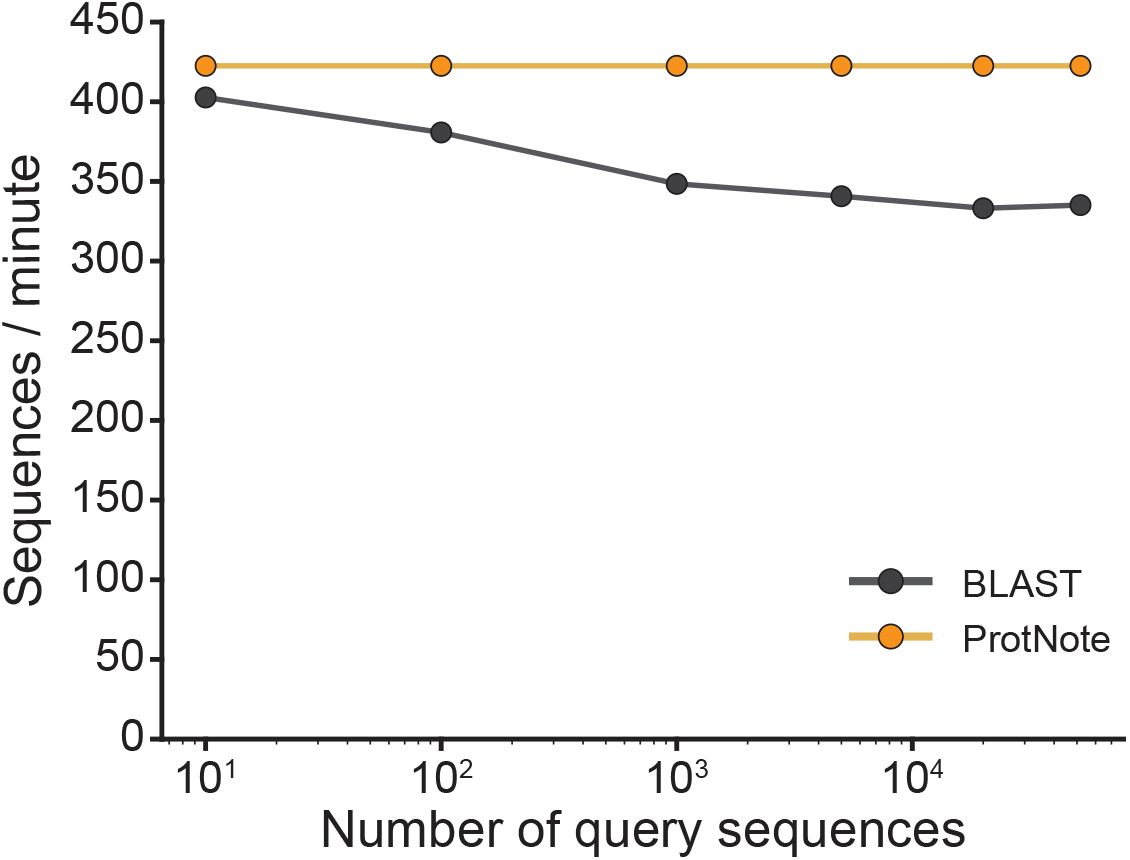
ProtNote is more efficient than BLAST at protein function prediction. Number of query sequences processed per minute (y-axis) for query sets of different sizes (x-axis) for ProtNote (yellow) vs BLAST (grey). ProtNote’s efficiency is constant as the query set increases, whereas BLAST’s efficiency decreases as the query set increases. Note that BLAST efficiency depends additionally on the training set size, which for these experiments was held constant.

**Fig. S10.**
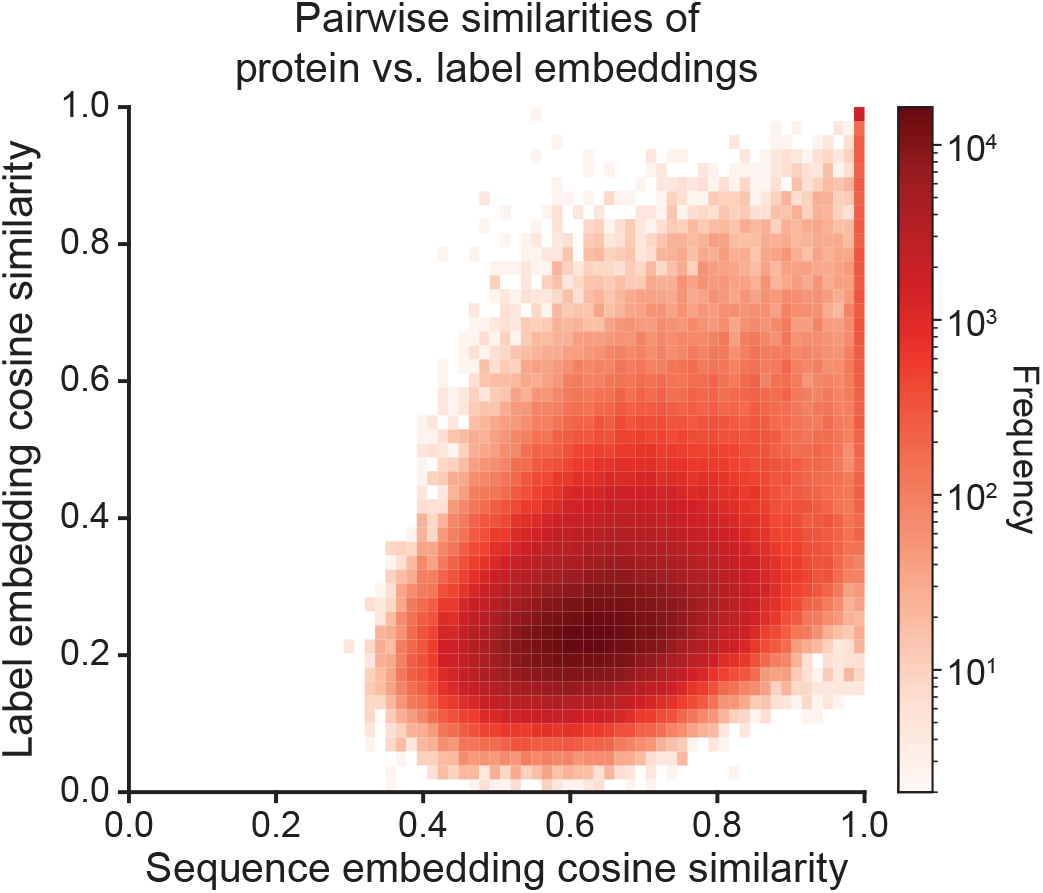
Correlation between pairwise similarities for protein sequence and label embeddings. All plots are based on a subset of the test set consisting of the most frequent GO annotations (*n*=1,502) for a random sample of sequences (*n*=800). Each label is represented in two ways: first, by the median sequence embedding of the sequences annotated with that label; second, with the label embedding from the label encoder projection head (before the final MLP). The plot shows a 2D histogram of the pairwise cosine similarities between the median sequence embeddings of the positive annotations (x-axis) versus the pairwise cosine similarities of the label embeddings (y-axis) for the label-label pairs of the sampled test set sequences (*n*=1502 median sequence embeddings x 1502 label embeddings = 2,256,004 pairs).

